# Local accessory gene sharing drives lineage-specific acquisition of antimicrobial resistance in Egyptian *Campylobacter spp*.

**DOI:** 10.1101/2021.09.24.461243

**Authors:** Shaimaa F. Mouftah, Ben Pascoe, Jessica K. Calland, Evangelos Mourkas, Naomi Tonkin, Charlotte Lefèvre, Danielle Deuker, Sunny Smith, Harry Wickenden, Matthew D. Hitchings, Samuel K. Sheppard, Mohamed Elhadidy

**Author notes:** Authors to which correspondence should be addressed. These authors contributed equally (Shaimaa F. Mouftah, Ben Pascoe). Division of Virology, Department of Pathology, University of Cambridge, Tennis Court Road, Cambridge, United Kingdom (Charlotte Lefèvre). Nuffield Department of Medicine, The Jenner Institute, University of Oxford, Oxford, United Kingdom (Danielle Deuker). **Repositories:** Short read data generated in this study are available on the NCBI Sequence Read Archive, associated with BioProject PRJNA576513 (https://www.ncbi.nlm.nih.gov/bioproject/PRJNA576513).

## Abstract

*Campylobacter* is the most common cause of bacterial gastroenteritis worldwide and diarrheal disease is a major cause of child morbidity, growth faltering and mortality in low- and middle-income countries (LMICs). Despite evidence of high incidence and differences in disease epidemiology, there is limited genomic data from studies in developing countries. In this study, we characterised the genetic diversity and accessory genome content of a collection of *Campylobacter* isolates from Cairo, Egypt. In total, 112 *Campylobacter* isolates were collected from broiler carcasses (n=31), milk and dairy products (n=24) and patients (n=57) suffering from gastroenteritis. Among the most common sequence types (STs) we identified were the globally disseminated, host generalist ST-21 clonal complex (CC21) and the poultry specialist CC206, CC464 and CC48. Notably, CC45 and the cattle-specialist CC42 were under-represented with a total absence of CC61. Comparative genomics were used to quantify core and accessory genome sharing among isolates from the same country compared to sharing between countries. Lineage-specific accessory genome sharing was significantly higher among isolates from the same country, particularly CC21 which demonstrated greater local geographical clustering. In contrast, no geographic clustering was noted in either the core or accessory genomes of the CC828, suggesting a highly admixed population. A greater proportion of *C. coli* isolates were multidrug resistant (MDR) compared to *C. jejuni*. This is a significant public health concern as MDR food chain pathogens are difficult to treat and often pose increased mortality risk demanding enhanced prevention strategies in the Egyptian market to combat such a threat.

**Impact statement:** *Campylobacter* is the leading bacterial cause of gastroenteritis worldwide and despite high incidence in low- and middle-income countries, where infection can be fatal, culture-based isolation is rare and the genotypes responsible for disease are seldom identified. Here, we sequenced the genomes of a collection of isolates from clinical cases and potential infection reservoirs from Cairo in Egypt and characterised their genetic diversity. Among the most common genotypes we identified were globally disseminated lineages implicated in human disease worldwide, including the host generalist ST-21 clonal complex (CC21) and the poultry specialist genotypes CC206, CC464 and CC48. Notably however, some other globally common genotypes were under-represented or entirely absent from our collection, including those from cattle-specialist lineages, CC42 and CC61. By focussing on specific lineages, we demonstrate that there is increased accessory genome sharing in specific clonal complexes. This increased local sharing of genes may have contributed to a greater proportion of *C. coli* isolates possessing antimicrobial resistance determinants that suggest they could be multidrug resistant (MDR). This is a significant public health concern as MDR food chain pathogens are difficult to treat and often pose increased mortality risk demanding enhanced prevention strategies.

**Data summary:** Short read data are available on the NCBI Sequence Read Archive, associated with BioProject PRJNA576513 (https://www.ncbi.nlm.nih.gov/bioproject/PRJNA576513). Assembled genomes, supplementary material and additional analysis files are available from FigShare: https://doi.org/10.6084/m9.figshare.9956597. Phylogenetic trees can be visualised and manipulated on Microreact for *C. jejuni* (https://next.microreact.org/project/Cjejuni_Egypt) and *C. coli* (https://next.microreact.org/project/Ccoli_Egypt) separately, or combined Cairo and Oxford data with additional PopPunk network clustering (https://microreact.org/project/Campy-Egypt).

## Introduction

Diarrheal disease is a major cause of child morbidity, growth faltering and mortality in low- and middle-income countries (LMICs) (McCormick and Lang, 2016; Platts-Mills and Kosek, 2014). *Campylobacter* is the most common cause of bacterial gastroenteritis worldwide (Kaakoush et al., 2015) and typically human campylobacteriosis is commonly diagnosed as a disease associated with consumption of contaminated food, especially poultry (Nichols et al., 2012; Sheppard et al., 2009). Extremely high incidence in LMICs, high exposure rates (Lee et al., 2013) and endemism among young children suggests a different epidemiology (Kaakoush et al., 2015; Lanata et al., 2013; J. Liu et al., 2016). Frequent or chronic (re)infection is allied to significant morbidity, cognitive development impairment, and even death (Coker, 2002; Crofts et al., 2018; Kirk et al., 2018; Reed et al., 1996). In Egypt, campylobacteriosis is common and a leading cause of paediatric diarrhoea, with an incidence of 1.2 episodes per year (ElGendy et al., 2018; Rao, 2001) with up to 85% of children infected in their first year (Liu et al., 2012). Despite the high frequency of reported cases of *Campylobacter*-associated diarrhoea in Egypt (ElGendy et al., 2018), there are no detailed surveillance studies on the dominant sequence types and proliferation of genotypes associated with the onset of post-infectious sequelae, such as irritable bowel syndrome (PI-IBS), Guillain-Barré syndrome (GBS) or Miller Fisher syndrome (Wierzba et al., 2008).

*Campylobacter* species are often part of the gut microbiota of various wild and farmed animals leading to frequent contamination of human food products (Asuming-Bediako et al., 2019; Waite and Taylor, 2015). In Egypt, farming practices can lack adequate biosecurity and regulation. Only limited studies have reported the prevalence and distribution of *Campylobacter* in Egyptian campylobacteriosis cases (Kaakoush et al., 2015) and little is known of the dominant source reservoirs driving infection and transmission. In Europe, potential source reservoirs have been identified through source attribution studies, with poultry products regarded as the primary source of infection (Facciolà et al., 2017; Mossong et al., 2016; Sheppard et al., 2009; Thépault et al., 2018). Host-adaptation of *Campylobacter* to a wide-range of hosts is reflected in its population structure (Colles and Maiden, 2012; Dearlove et al., 2016; Griekspoor et al., 2013; Méric et al., 2018; Sheppard et al., 2014), with many lineages common in human infection able to infect multiple host species. These host generalist lineages include *C. jejuni* ST-21, ST-45 clonal complexes and the *C. coli* ST-828 complex (Dearlove et al., 2016; Mossong et al., 2016). Other genotypes are only found in a single reservoir species, often associated with global poultry or cattle production. Host specialist clonal complexes common in human disease includes the poultry-associated ST-353, ST354 and ST257 (Berthenet et al., 2019; Sheppard et al., 2009) and cattle specialist ST-61 (French et al., 2005; Mourkas et al., 2019).

Human infection in developed countries is usually sporadic and self-limiting, not requiring treatment with antibiotics. However global rates of antimicrobial resistance are rising (Mourkas et al., 2019; Zhao et al., 2016) in line with other Gram negative gastrointestinal pathogens (Tam et al., 2012; CDC, 2020). Widespread agricultural usage has driven the proliferation of tetracycline resistance through its use as a growth promoter (Abdi Hachesoo et al., 2014; Inglis et al., 2019). In particular, *C. coli* has shown an ability to acquire erythromycin resistance genes from other species (Mourkas et al., 2019). This has not been explored for Egyptian *Campylobacter* isolates, where agricultural antibiotic usage is poorly regulated (Dahshan et al., 2015) and self-medication for gastrointestinal disease is common (Abd El-Tawab et al., 2018; Sabry et al., 2014). Global differences in the use of quinolones is likely responsible for the geographical differences observed in quinolone resistance (Luangtongkum et al., 2009; Pascoe et al., 2017; Zollner-Schwetz and Krause, 2015).

We have sequenced 112 *Campylobacter* isolates collected from patients and food of animal source (i.e., broiler chicken carcasses and dairy products) in Cairo over a year to determine the most prevalent *Campylobacter* genotypes causing disease in Egypt. By screening the genome content, including known AMR determinants we provide a better understanding of the local population structure to guide disease intervention in Egypt. This study provides a basis for considering complex transmission networks in LMICs and highlights the role of globally transmitted *Campylobacter* lineages and the emergence of (horizontally acquired) antimicrobial resistance.

## Methods

### Ethical approval

The study represents a retrospective study that involved sequencing the genomes of a historical strain collection and no patient data collection was involved in this study. Ethical approval was granted from the respective ethics committee in the Egyptian central directorate of research and health development before conducting the study.

### Isolate collection

In total, 112 *Campylobacter* isolates were collected in Cairo, Egypt from September 2017 to December 2018, including 31 isolates from broiler carcasses, 24 isolates from milk and dairy products, and 57 clinical isolates. Clinical isolates were recovered from stool samples of patients admitted to hospitals in downtown Cairo suffering from gastroenteritis symptoms. A questionnaire was distributed to all admitted patients requesting details on clinical presentation (e.g., duration of illness, symptoms, medication prescribed), dietary record of the previous 2 weeks, including consumption of specific or undercooked meats, unpasteurized milk, exposure to animal manure or faeces, and any retail outlets commonly used by patients for food consumption prior to the onset of illness. A random sampling approach was then used to include food samples from stores in the study region that were commonly listed in the questionnaire.

### Sample culturing and whole genome sequencing

The isolation and enumeration of *Campylobacter* strains from different food matrices was performed according to the ISO 10272-1 (Enrichment Method; Detection of *Campylobacter* spp. after Selective Enrichment). All isolates were sub-cultured from -80°C frozen stocks onto Mueller-Hinton agar (Oxoid, United Kingdom). Plates were incubated at 42 ± 1°C under anaerobic conditions using AnaeroGen™ 2.5L Sachets (Oxoid, United Kingdom). Genomic DNA was extracted from 112 Egyptian isolates using the QIAamp DNA Mini Kit (QIAGEN, Crawley, UK), according to manufacturer’s instructions and DNA concentrations were quantified using a Nanodrop spectrophotometer before genome sequencing using an Illumina MiSeq (California, USA). Nextera XT libraries (Illumina, California, USA) were prepared following manufacturer’s protocols and short paired-end reads were sequenced using 2□× □300□bp paired end v3 reagent kit (Illumina).

### Genome datasets

Genomes were assembled *de novo* using SPAdes (version 3.8.0; Bankevich et al. 2012). The average number of contigs was 72 (range: 12–471) for an average total assembled sequence size of 1.70 Mbp (range: 1.56–1.86). The average N50 contig length (L50) was 14,577 (range: 3,794-55,912) and the average GC content was 30.8 % (range: 30.5-31.6). Short read data are available on the NCBI short read archive (SRA), associated with BioProject PRJNA576513. Assembled genomes and supplementary material are available from FigShare (doi:10.6084/m9.figshare.9956597; individual accession numbers and assembled genome statistics in **Supplementary Table S1**). We augmented our collection by assembling a context dataset of previously published isolates (n=204) to represent the known diversity of *C. jejuni* and *C. coli* (Calland et al., 2020; Sheppard et al., 2010, 2013, 2014). In addition, we also compared our single city survey with a previously published survey from Oxford in the UK (n=874 isolates collected over 1 year; Cody et al. 2012). Isolate genomes were archived in BIGSdb and MLST sequence types (STs) derived through BLAST comparison with the pubMLST database (Dingle et al., 2001; Jolley et al., 2018; Jolley and Maiden, 2010; Sheppard et al., 2012). Simpson’s index of ST diversity was calculated for the Cairo and Oxford datasets using the equation:

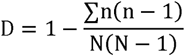

Where *n* is the number of isolates of each sequence type and *N* is the total number of isolates (Grundmann et al., 2001).

### Core and accessory genome characterisation

Alignments were made from concatenated gene sequences of all core genes (found in ≥95% isolates) using MAFFT (version 7; Katoh and Standley 2013) on a gene-by-gene basis. Separate maximum-likelihood phylogenies were constructed with a GTR+I+G substitution model and ultra-fast bootstrapping (1000 bootstraps) (Hoang et al., 2018) implemented in IQ-TREE (version 1.6.8; Nguyen et al. 2015) for *C. jejuni* (n=1,048) and *C. coli* (n=132) and visualized on Microreact (https://next.microreact.org/project/Cjejuni_Egypt; https://next.microreact.org/project/Ccoli_Egypt) (Argimón et al., 2016).

All unique genes present in at least one isolate (the pangenome) were identified by automated annotation using PROKKA (version 1.13; Seemann 2014) followed by PIRATE, a pangenomics tool that allows for orthologue gene clustering in bacteria (Bayliss et al., 2019). We defined genes in PIRATE using a wide range of amino acid percentage sequence identity thresholds for Markov Cluster algorithm (MCL) clustering (45, 50, 60, 70, 80, 90, 95, 98). Genes in the pangenome were ordered initially using the NCTC 11168 reference followed by the order defined in PIRATE based on gene synteny and frequency (Gundogdu et al., 2007; Pascoe et al., 2019). As described previously, a matrix was produced summarizing the presence/absence and allelic diversity of every gene in the pangenome list, with core genes defined as present in 95% of the genomes and accessory genes as present in at least one isolate (**Supplementary table S2**) (Méric et al., 2014). Pairwise core and accessory genome distances were compared using PopPunk (version 2.2.0; Lees et al. 2019) which uses pairwise nucleotide k-mer comparisons to distinguish shared sequence and gene content to identify divergence of the accessory genome in relation to the core genome. A two-component Gaussian mixture model was used to construct a network to define clusters, comparable to other *Campylobacter* studies (Components: 41; density 0.0579; transitivity: 0.9518; score: 0.8907) (Pascoe et al. 2020).

Core genome variation between isolates was quantified by calculating the pairwise average nucleotide identity (ANI) of all (n=112+874) *Campylobacter* genomes using FastANI v.1.058 (Jain et al., 2018). The gene presence matrix produced by PIRATE was used to generate a heatmap of shared pairwise accessory genome genes. Averages were calculated for within and between country comparisons in addition to focussed analysis on the ST21 (*C. jejuni*) and ST828 (*C. coli*) clonal complexes. Antimicrobial resistance genes and putative virulence genes were detected through comparison with reference nucleotide sequences using ABRicate (version 0.8) (https://github.com/tseemann/abricate) and the NCBI database (Chen et al., 2005; NCBI Resource Coordinators, 2013). Point mutations related to antibiotic resistance genes were identified by PointFinder (Zankari et al., 2017) using the STAR-AMR software package (https://github.com/phac-nml/staramr) (**Supplementary table S3**).

## Results

### Globally circulating genotypes among Egyptian Campylobacter isolates

We sequenced and characterized a collection of *Campylobacter spp*. isolates (n=112) from clinical cases, broiler carcasses and dairy products collected over a 14-month sampling period in Cairo, Egypt (**Figure 1A; Supplementary table S1**). Isolate genotypes were compared with all genomes deposited in the pubMLST database (97,012 profiles, data accessed 17^th^ February 2020) and ranked according to how frequently they were found associated with human disease (**Figure 1B**). Egyptian *C. jejuni* isolates belonged to 15 clonal complexes (CCs) with a diverse assemblage of STs. Nearly half of the isolates (n = 29, 47%) were from common lineages, isolated many times before and recorded in pubMLST (>50 MLST profiles; **Figure 1B**), including the globally disseminated lineages of ST-21CC (n=37; 41%), ST-206 CC (n=10; 11%) and ST-464CC (n=7; 8%) the most abundant. Several other poultry-associated clonal complexes, which are common in human disease (Berthenet et al., 2019; Sheppard et al., 2009), including ST-353 (n□=3, □3.2%,), ST-354 (n□= □4, 4.3%) and ST-257 (n□= □4, 4.3%) were identified. Other globally disseminated lineages were found less often in Egypt (n<=2) i.e., ST□460 (n□= □2), ST□1034 (n□= □2), ST□42 (n□= □1), ST-45 CC (n=1), ST□573 (n□= □1), ST□574 (n=1) and ST□658 (n=1) (Colles et al., 2010; Olkkola et al., 2016).

**Figure 1:**
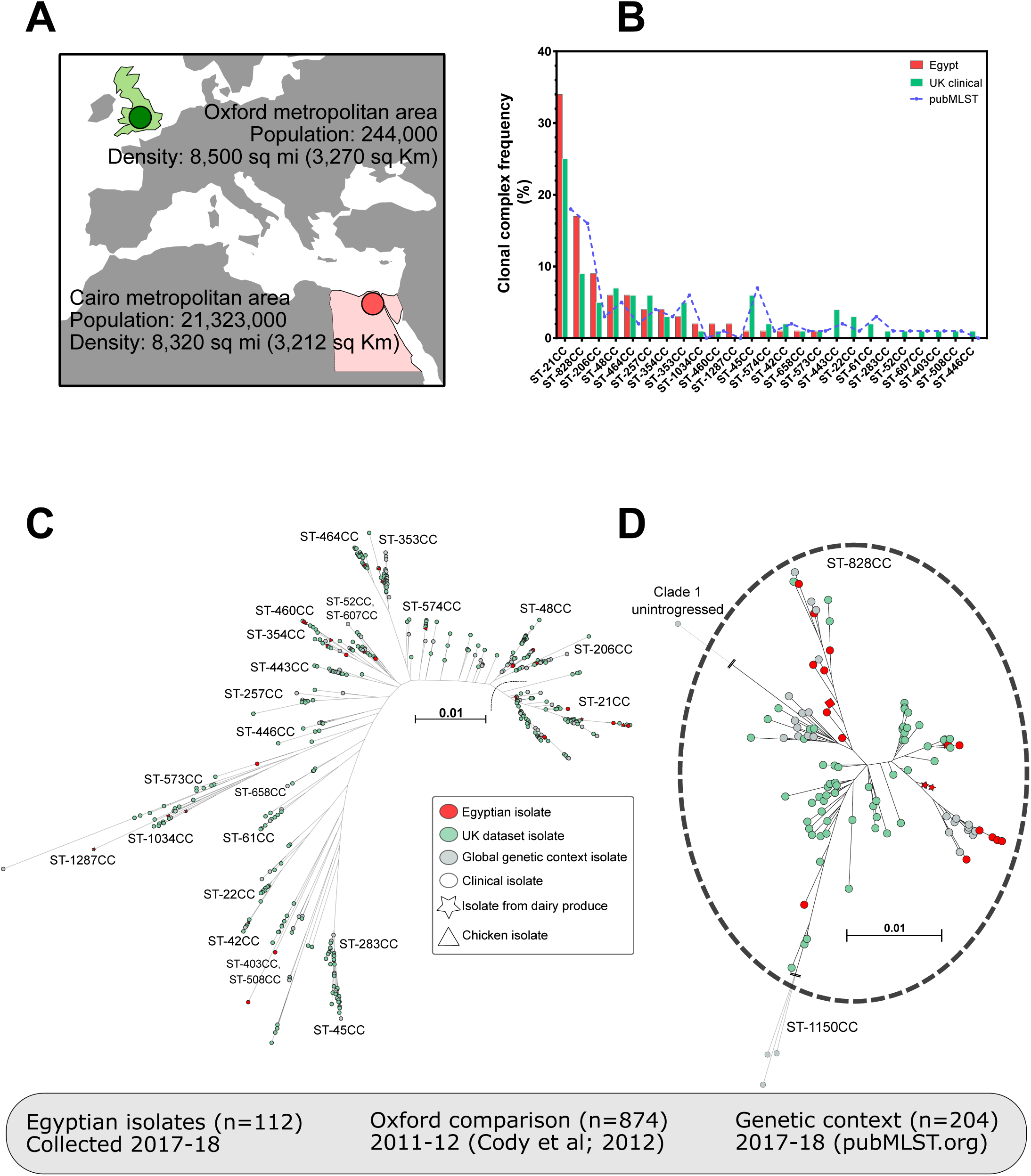
**(A)** Demographic data for Cairo, Egypt from which we collected *Campylobacter spp*. isolates (n=112; red circles) from clinical cases, broiler carcasses and dairy products collected over a 14-month sampling period. Our collection was compared to a similar published survey from Oxford, UK (n=874; green circles; Cody et al. 2012) and isolates from pubMLST.org (n=204; grey circles) for additional genetic context. (**B)** Clonal complexes (CCs) of isolates collected from Cairo were ranked according to the frequency in our local dataset and how often they have been sampled from human disease isolates (data from pubMLST; https://pubmlst.org/). Alignments were made from concatenated gene sequences of all core genes (found in ≥95% isolates) using MAFFT (version 7; Katoh and Standley 2013) on a gene-by-gene basis. Separate maximum-likelihood phylogenies were constructed with a GTR+I+G substitution model and ultra-fast bootstrapping (1000 bootstraps) (Hoang et al., 2018) implemented in IQ-TREE (version 1.6.8; Nguyen et al. 2015) for (**C**) *C. jejuni* (n=1,048) and (**D**) *C. coli* (n=132) and visualized on Microreact (https://next.microreact.org/project/Cjejuni_Egypt; https://next.microreact.org/project/Ccoli_Egypt) (Argimón et al., 2016).

### Local sequence types

Comparison with a collection representing the known genetic diversity of *C. jejuni* and *C. coli* identified some common STs (>1,000 profiles in pubMLST) that were completely absent in our Egyptian collection, i.e., ST-53, ST-829 (*C. coli*), ST-22, ST-61, ST-51, ST-1068 (*C. jejuni*) (**Figure 1CD)**. Two isolates belonging to ST-1287CC, a genotype that has previously been isolated from poultry and the environment (Magnússon et al., 2011), was observed exclusively among our Egyptian isolates, yet absent in UK and genetic context datasets. Furthermore, there were also some STs belonging to ST-21CC that were found in Egyptian isolate collection (n=>3) that are rare in global collections (<100 profiles in pubMLST), i.e., ST-1519 (n=4), ST-3769 (n=3). It was also observed that more *C. coli* was found among Egyptian clinical isolates than is typically observed, specifically the *C. coli* lineage ST-828 CC 90.4% *C. coli* isolates (19/21) belonged to the ST-828 CC within the Egyptian dataset and two *C. coli* isolates with unassigned CC of sequence types, ST-7951 and ST-1681. Three rare STs belonging to ST-828 CC were exclusively found in Egypt dataset which are ST-1058 (n=1), ST-1059 (n=1), and ST-7950 (n=1).

### Increased sharing of accessory genes contributes to a local gene pool

Our Egyptian dataset was compared directly with a previously published study of a single city, ∼1-year survey from Oxford in the UK (Cody et al., 2012). Both populations were similarly diverse, specifically there were 50 STs (16 CCs) among the Egyptian isolate collection, with a Simpson’s diversity index of 0.817, compared to 205 STs (32 CCs) among the Oxford collection of genomes (Simpson’s diversity index = 0.895; **Figure 1CD**). We used PIRATE to construct a pan-genome of all Egyptian and Oxford isolates (n=986). Consistent with other studies, we identified an open pangenome, meaning that the number of genes in the pangenome continues to increase with each additionally sequenced isolate. Accessory genes represented nearly three-quarters of the pangenome (3,410 genes; 74% of pangenome) with a quarter of the genes identified (1,225, 26%) considered core genes present in 95% or more of the isolates. Pairwise comparison of the core nucleotide sequence (%ANI) and accessory genome sharing of all isolates reflected the clonal frame, with clusters of closely related isolates sharing a large percentage of ANI (**Figure 2AB**). Direct comparison between the Oxford and Cairo datasets suggested an increase in within-country, local accessory gene sharing (**Figure 2CD**). The structured clustering of pairwise comparisons of shared accessory genes suggested that this may vary between lineages and visualization of the differences in the distribution of pairwise genomic distances with PopPUNK also pointed towards lineage-specific shared gene pools (**Figure 2E**). Host generalist clonal complex isolates clustered closer together than the more isolated host-specific isolates. This included the two most common clonal complexes identified in our Cairo collection, ST-21CC and ST-828CC, which were investigated further (**Figure 2F**).

**Figure 2:**
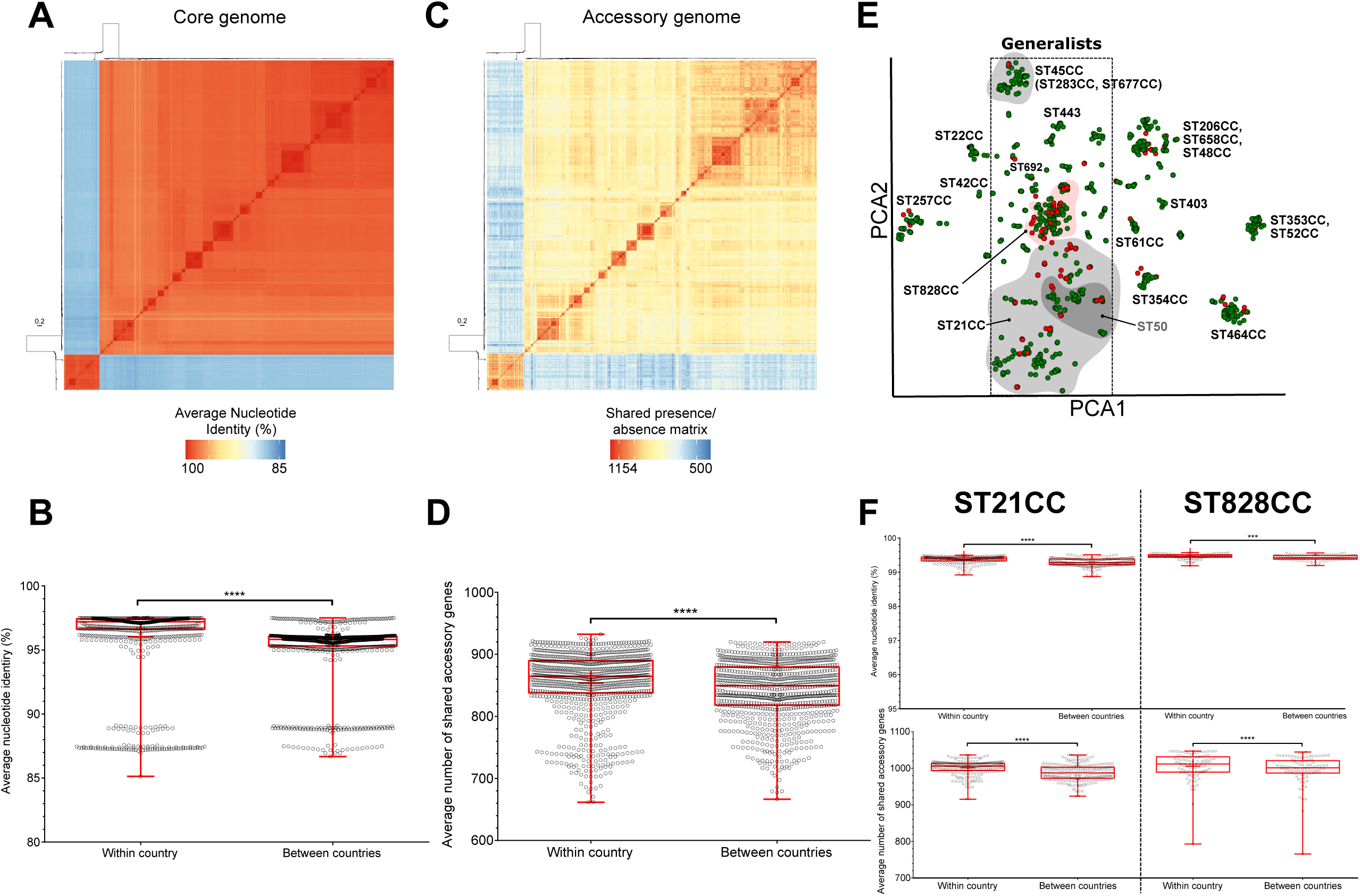
**(A)** Core genome variation between isolates was quantified by calculating the pairwise average nucleotide identity (ANI) of all UK and Oxford *Campylobacter* genomes (n=112+874) using FastANI v.1.058 (Jain et al., 2018). **(B)** The ANI for each isolate was estimated and averages compared within and between countries. **(C)** The gene presence matrix produced by PIRATE was used to generate a heatmap of shared pairwise accessory genome genes. **(D)** Averages were calculated for within and between country. **(E)** Clustering of pairwise core and accessory genome distances were compared using PopPunk. Interactive visualisation on Microreact: https://microreact.org/project/Campy-Egypt. **(F)** Comparisons of within and between country ANI and accessory gene sharing were also analysed for our two most common Egyptian lineages, ST21 (*C. jejuni*) and ST828 (*C. coli*) clonal complexes.

### Locally diverged sequence types within the globally disseminated ST-21 clonal complexes

As one might expect of within lineage (clonal complex) comparisons, all ST-21CC isolates shared more than 99% core genome nucleotide identity and shared more accessory genes than the population average (852 genes; **Figure 2CF**) and significantly more genes were shared between isolates from the same country (*t-*test with Welch correction; p<0.0001). A maximum-likelihood phylogeny of all CC21 isolates (n=251), the most common clonal complex identified in our collection from Cairo, identified geography-specific clusters of isolates (**Figure 3A**). These clones also clustered together when visualizing the distribution of pairwise genomic distances with PopPUNK (**Figure 3B**). While some specific STs were common in both Oxford and Cairo (ST21 and ST50), others were much more common in one specific location, e.g., ST-53 in Oxford, and ST-1519 and ST-3769 in Cairo (**Figure 3C**). There was also evidence that some lineages had enhanced AMR (**Figure 3D**). While the ST-50 genotype is very common and has been reported more than 3,900 times in pubMLST from 40 countries, this among the first reports from Africa. In both Oxford and Cairo datasets, ST-50 was often predicted to be MDR. ST-21 is also very common, with more than 4,000 reports from 33 countries in pubMLST but was much less likely to be MDR. Four isolates of the Cairo specific ST-3769 also represented a high proportion of MDR (**Figure 3E**).

**Figure 3:**
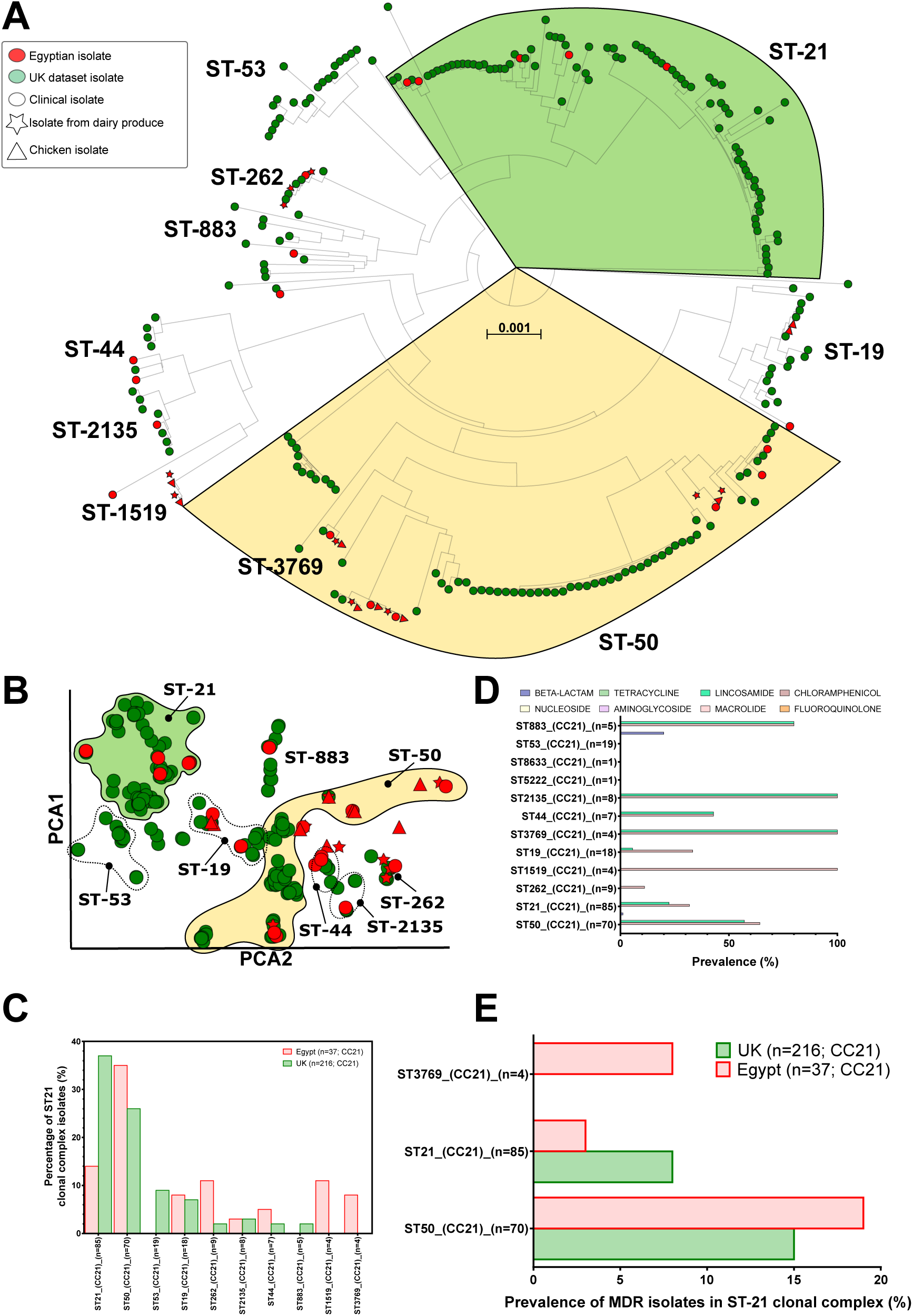
**(A)** Sub-tree of all Egyptian and UK ST21 clonal complex (CC21) isolates (n=251). Common sequence types are annotated and ST50 (yellow) and ST21 (green) are highlighted. **(B)** Within clonal complex clustering of pairwise core and accessory genome distances with PopPunk. **(C)** Prevalence of the most common sequence types found within CC21. **(D)** Prevalence of AMR determinants grouped by antibiotic class for each CC21 ST. **(E)** Prevalence of MDR isolates (AMR determinants for three or more antibiotic classes) in CC21 STs.

### Extensive multi-drug resistance in local C. coli sequence types

Greater admixture was noted between UK and Egyptian ST-828CC isolates than for ST-21CC – no geographic clustering was observed in either the core or accessory genomes (**Figure 4AB**). However, only ST-827 was common in both datasets (**Figure 4C**). Several STs were found in the Oxford dataset that were not identified in Cairo, including the frequently isolated STs -829, -828, -855, 962, -1145 and -5734. Several lineages were highly resistant to lincosamides, with more than half the isolates from ST-828, ST-830 and ST-872 predicted to be resistant (**Figure 4D**). All isolates from ST828 and ST-872 were also predicted to be resistant to chloramphenicol. Overall, *C. coli* isolates (6 of 105, 5.7%) were far more likely to be considered MDR than *C. jejuni* isolates (6 of 876, 0.68%) and ST-828 complex isolates from Cairo (2 of 19, 10.5%) demonstrated much higher rates of MDR than in Oxford (3 of 77, 3.8%; **Figure 4E**).

**Figure 4:**
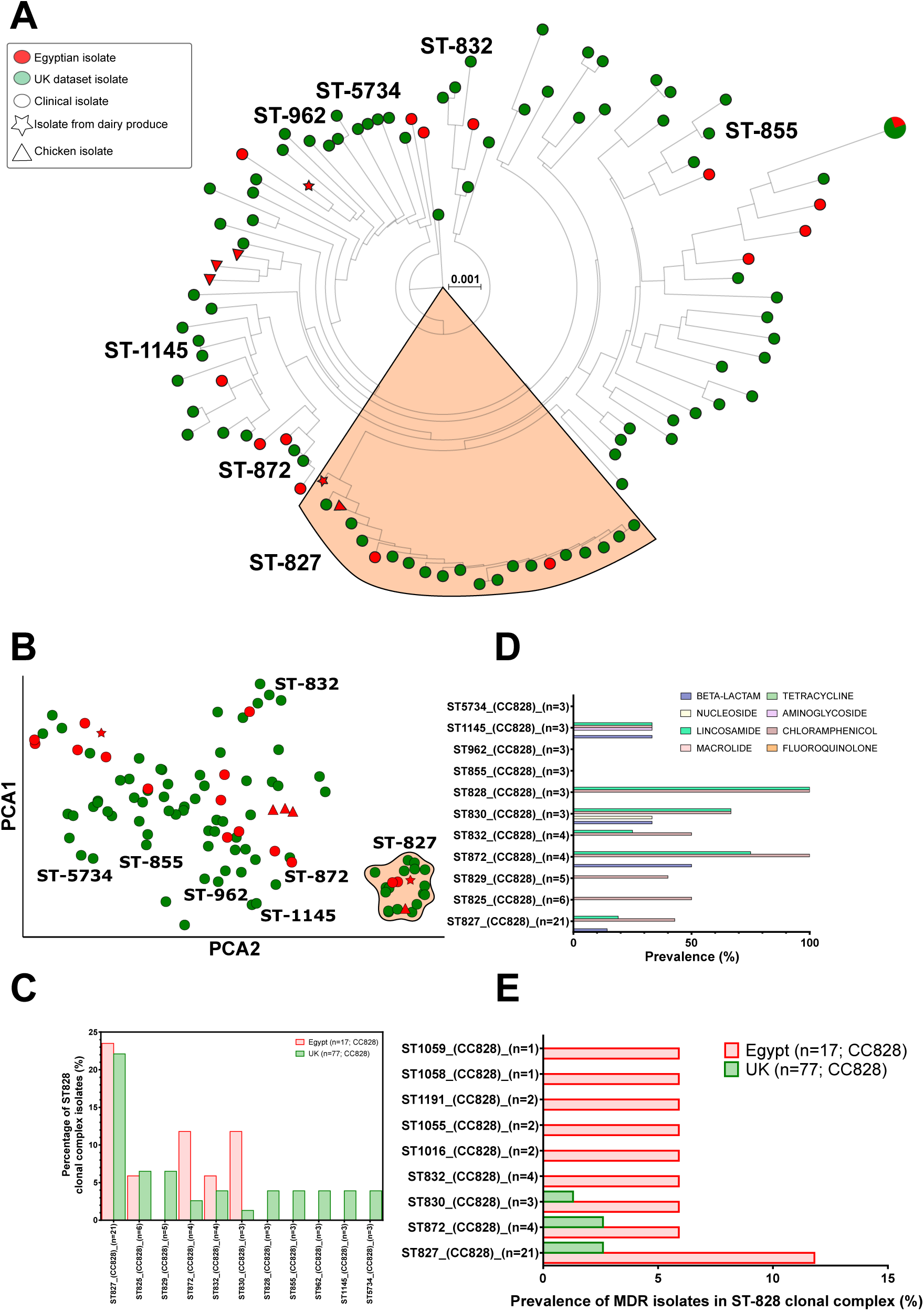
**(A)** Sub-tree of all Egyptian and UK ST828 clonal complex (CC828) isolates (n=94). Common sequence types are annotated and ST827 (orange) is highlighted. **(B)** Within clonal complex clustering of pairwise core and accessory genome distances with PopPunk. **(C)** Prevalence of the most common sequence types found within CC828. **(D)** Prevalence of AMR determinants grouped by antibiotic class for each CC828 ST. **(E)** Prevalence of MDR isolates (AMR determinants for three or more antibiotic classes) in CC828 STs.

### Antimicrobial resistance genes are distributed across isolates

In characterization of the resistome, each isolate genome was screened for the presence of genes associated with AMR. In Egypt, for *C. jejuni*, the average number of AMR genes per isolate was 6.66, comparable to 6.52 for *C. coli*. In Egypt, the presence of the *tet(O)* gene, conferring tetracycline resistance, was higher in *C. coli* than *C. jejuni* (76% and 43% respectively). This pattern contrasts with Oxford where 41.7% of *C. jejuni* but only 35.3% of *C. coli* isolates were found to harbor *tet(O)*. Whilst a low proportion of Egyptian isolates (6.7%) contained the *blaOXA-61* gene, associated with β-lactam resistance, alternative alleles including *blaOXA-450* and *blaOXA-605* were abundant. In respect to lineage association with genes, in Egypt the ST-21 clonal complex had a high prevalence of genes associated with β-lactam resistance (particularly the blaOXA-193, blaOXA-450 and blaOXA-605 alleles). The blaOXA-465 allele was closely related to ST-1034. Furthermore, blaOXA-61 was closely associated with ST-48 (**Figure 3D**). All of these patterns were reflected amongst the Oxford isolates. However, numerous genes (*including aadE, Ant6-la* and *blaOXA-451*) were found amongst distant lineages. The multi-drug efflux pump encoded by a three-gene operon (cmeABC) was abundant amongst isolates (n=87,74%) – although an absence of the repressor gene cmeR in *C. coli* was observed.

Whilst the average number of resistance genes per isolate was comparable for *C. jejuni* in Egypt, this analysis indicated that *C. coli* held a greater breadth of genes across classes of antimicrobials. Hence, the proportion of MDR isolates, considered when an isolate is resistant to at least three classes, was 28% for *C. coli* compared to 1% for *C. jejuni* (EFSA, 2021). The majority (88%) of MDR isolates in Egypt were *C. coli*, despite *C. coli* representing about a fifth of the dataset. In other words, a greater proportion of *C. coli* isolates were MDR. In Oxford, half of MDR isolates were *C. coli*, whilst in this case representing less than one tenth of the dataset. The *C. jejuni* isolates that were MDR, were all host generalists – ST-21, ST-48 or ST-206. Amongst Egyptian isolates, genes including *aad9, aadE, aadE-Cc, ant(6)-Ia, aph(2’’)-If, aph(3’)-III* and *aph(3’)-IIIa* associated with aminoglycoside resistance, were almost exclusively associated with *C. coli*, particularly MDR *C. coli*. This association was not as strong in Oxford. Regarding specific genes and host associations, aminoglycoside resistance-associated genes were infrequent amongst isolates from chicken or dairy products. *ant(6)-Ia* for example, was solely found in human samples. In turn, few isolates from chicken and dairy products were MDR (only 12.5% of MDR isolates was from chicken).

## Discussion

Diarrheal disease is a major threat to human health and the second leading cause of death in children under five years’ old LMICs (Lanata et al., 2013). Campylobacteriosis is a major cause of diarrheal disease worldwide (Amour et al., 2016; ElGendy et al., 2018; Lee et al., 2013) but, despite the potential importance, little is known about *Campylobacter* in countries where it potentially poses the greatest health risk. As studies begin to take a worldview of *Campylobacter* epidemiology and transmission (Mottet and Tempio, 2017), we describe globally disseminated agriculture-associated disease-causing lineages based on core and accessory genome content, with evidence that local accessory genome sharing driving acquisition of AMR genes in specific lineages.

The Egyptian *Campylobacter* isolates included a diverse set of STs, including common disease-causing lineages and regional STs, that have rarely been reported from other parts of the world. Industrialized agriculture globalization has dispersed livestock worldwide (Mottet and Tempio, 2017), expanding the geographical range of *C. jejuni*. This is evident in the Egyptian collection as two of the most predominant genotypes belonged to the ST-21 and ST-206 clonal complexes (Figure 1D). These two host generalist clonal complexes have been extensively reported worldwide and frequently isolated from various reservoir hosts, including human clinical samples (Berthenet et al., 2019; Dingle et al., 2001; Grove-White et al., 2011; Mossong et al., 2016; Sheppard et al., 2009; Suerbaum et al., 2001). The ST21-CC exhibits considerable genome plasticity with a clear association with several virulence genes and resistance to various antimicrobial agents (Aksomaitiene et al., 2019; Gripp et al., 2011; Habib et al., 2010; Wieczorek et al., 2017; T. Zhang et al., 2016). Poultry-associated clonal complexes, ST-206, ST-464, ST-48, ST-257 and ST-354 were also common among the Egyptian isolates, all of which are among the most prevalent clonal complexes isolated in Europe (Colles et al., 2011; Elhadidy et al., 2018; Fiedoruk et al., 2019).

Further comparison of isolate genotypes collected in Cairo with a large global collection revealed the absence of certain lineages, most notably the lack of the cattle-associated genotype, ST-61 (Dingle et al., 2002; Mourkas et al., 2020). There was only one isolate, of dairy product origin, that could be attributed to a cattle-specialist clonal complex (ST-42), which is unexpected as several (n=24) isolates were sampled from dairy products. *Campylobacter* isolates from cattle have predominantly been sampled from meat, milk products and fecal sources (n=2,726 in pubMLST; Kwan et al., 2008; Mourkas et al., 2020; Epping et al., 2021). Suggesting that dairy products isolates might represent a different source population in Egypt.

There were also no isolates belonging to the ST-22 CC, a particularly high risk lineage which is commonly found among patients with post-infectious complications of campylobacteriosis, such as GBS and IBS (Revez et al., 2011; Peters et al., 2021). Although one isolate in our collection was from ST-45 CC, this host generalist clonal complex is often one of the most commonly isolates lineages in clinical surveillance studies worldwide (De Haan et al., 2010; Sheppard et al., 2009; Shin et al., 2013; Sopwith et al., 2008). Notably however, it is often absent (or under-represented) in studies conducted in LMICs (Pascoe et al., 2020; Sarhangi et al., 2021). This is consistent with observations from other LMICs, where local differences in disease epidemiology are reflected by the absence of common *Campylobacter* lineages, and the presence of rare or unique sequence types (Graham et al., 2016; Pascoe et al., 2020; Prachantasena et al., 2016; P. Zhang et al., 2020). Among our Egyptian isolates the ST-1287 clonal complex (n=2) has been reported less than 4 times from other parts of the world (Colles et al., 2011; de Haan et al., 2010; Ramonaite et al., 2014; P. Zhang et al., 2020).

Geographical differences have been noted in ST-21CC (Kärenlampi et al., 2007; Kovanen et al., 2014; Olkkola et al., 2016; Pascoe et al., 2017; Wallace et al., 2021). ST-21 CC isolates are among the most common *C. jejuni* genotypes isolated worldwide, with one quarter of *C. jejuni* isolates recorded in the pubMLST database are ST21 CC. Isolates of the ST-50 sequence type (n= 3,915) alone have been sampled from 6 continents and 44 countries, although this will be their first report from Africa (Jolley et al., 2018). Our Egyptian ST-50 isolates do cluster together on a ML phylogeny of ST-21 CC isolates and away from the Oxford ST-21 CC when grouped by PopPunk. Two sequence types were unique to Egypt, ST-1519 and ST-3769, with nearly 10% of the ST-3769 isolates were MDR. A slightly greater proportion of the Egyptian ST-50 isolates were also MDR, although this sequence type has been observed to be MDR in other parts of the world (Elhadidy et al., 2020).

The *C. coli* ST-828 clonal complex did not show as much geographical segregation, and when grouping our Egyptian isolates by core and accessory genome distances they clustered with the UK isolates, despite several STs being isolated in only one of the datasets. STs found in the Egyptian dataset were more often MDR than UK isolates, and overall *C. coli* from Cairo were far more MDR than *C. coli* isolates from developed countries (Du et al., 2018; Gharbi et al., 2018; Mourkas et al., 2019). The most compelling clarification for such abundance could be that *C. coli* of ST-828 CC have a great recombination potential besides the accumulation of *C. jejuni* DNA throughout the genome of this lineage which could have led to the acquisition of multiple AMR genes (Sheppard et al., 2008, 2013).

Overall, there is a clear evidence of local sharing and recent acquisition of accessory gene content of AMR genes within the Egyptian isolates. Specifically, pairwise clustering of isolates by core and accessory genome distances recapitulated clusters according to ST and clonal complex (**Figure 2**), however most Egyptian isolates were more tightly clustered than the Oxford dataset, consistent with shared acquisition of accessory genes. Overall, ANI and shared accessory genes were similar between Oxford and Egyptian isolates (per isolate), however the two most common clonal complexes found in our Cairo dataset demonstrated greater sharing of accessory genes, indicative of a shared gene pool. Our study suggested that while geographical partitioning doesn’t impact the composition of the core genome, represented by the shared STs and CCs, the accessory genome is influenced. Within the Egyptian isolates, the most prevalent *C. jejuni* genotypes (ST-21CC and ST-206CC) showed clear evidence of transmission of MDR determinants among lineages. Multiple factors could influence this, such as livestock and food production practices and the segregation of MDR isoaltes. However, selective pressure for MDR is clearly attributable to antibiotic usage and potentially zoonotic transmissions as well as the rate of horizontal gene transfer (Fiedoruk et al., 2019). Our study provides evidence to support programs aimed at improved antibiotic stewardship in clinical and veterinary settings. With strict control measures, and an understanding of transmission of strains from animal reservoirs through the food production chain, it may be possible to reduce contamination with MDR *Campylobacter* in Egypt.

## Author statements

### Author contributions

SM, JKC, BP, ME and SKS designed the study and wrote the paper.

JKC, BP, EM, GF, CL, HW performed genomic analysis.

BP, CL and MDH sequenced and assembled genomes.

All authors contributed and approved the final manuscript.

## Supplementary information

**Supplementary table 1:** Summary of isolate collection data and genome statistics

**Supplementary table 2:** Summary PIRATE core and accessory genome statistics.

**Supplementary table 3:** Summary of AMR genes identified by comparison with the NCBI database and pointfinder.

